# Csmd2 interacts with Dab1 and is Required in Reelin-Mediated Neuronal Maturation

**DOI:** 10.1101/2020.01.30.925537

**Authors:** Mark A Gutierrez, Brett E Dwyer, Santos J Franco

## Abstract

Reelin is a glycoprotein secreted by Cajal-Retzius cells to regulate development of the cerebral cortex. Reelin binding to its receptors on immature neurons initiates a signaling cascade through the downstream adaptor protein, Dab1. Defects in this signaling mechanism result in perturbed neuronal migration, reductions in dendrite complexity, and deficits in synapse development and function. How Reelin controls neuronal migration and brain lamination have been extensively investigated over the years, but the pathways that regulate dendrite and spine development downstream of Reelin and Dab1 have yet to be fully elucidated. Here, we have identified a novel interaction between Dab1 and Csmd2, a synaptic transmembrane protein required for dendrite and dendritic spine development in forebrain excitatory neurons. We demonstrate that Csmd2 contains an NPxY motif on its intracellular region, through which Dab1 interacts with Csmd2. Interestingly, we find that this NPxY consensus motif is not required for Csmd2 to localize at the postsynaptic densities of spiny neurons. Rather, the introduction of an NPxY mutant form of Csmd2 results in a significant overproduction of immature, filopodia-like dendritic spines in maturing neurons. Moreover, we show that knockdown of *Csmd2* mRNA expression in immature developing neurons abolishes the ability of Reelin to promote dendrite elaboration and dendritic spine maturation. This suggests that the Csmd2-Dab1 interaction may be a requirement of Reelin/Dab1 signaling to mediate the structural maturation of neurons. Together, these results point toward a role of Csmd2 in the Reelin/Dab1 signaling axis that promotes the development of dendrites and dendritic spines in maturing neurons.

**Summary Statement:** How Reelin controls neuronal maturation remains to be understood. We demonstrate that the synaptic protein Csmd2 interacts with the Reelin-associated adaptor protein Dab1. We also determine that Reelin requires Csmd2 to regulate structural development and maturation of forebrain neurons.

## Introduction

During development of the mammalian forebrain, a series of cellular migration events organize neurons into specific cell layers prior to the projection and elaboration of dendritic arbors and the establishment of functional neural circuits through synaptic connections. One of the most well-known factors required for these aspects of forebrain development is the secreted glycoprotein Reelin. The Reelin loss-of-function (*reeler*) mutant was first reported in 1951 (Falconer, 1951), the phenotype of which was later described as widespread cellular disorganization and lamination defects (Rice et al., 1998; Olson et al., 2006; Franco et al., 2011). Such a mutation has been reported to be associated with the risk of neuropsychiatric disorders, including schizophrenia and autism spectrum disorder (Eastwood and Harrison, 2006; Ovadia and Shifman, 2011; Lammert et al., 2017; Sanchez-Sanchez et al., 2018; Sobue et al., 2018). Therefore, identifying and understanding the function of the molecular factors through which Reelin governs brain development is an endeavor of significant clinical relevance.

Upon secretion by Cajal-Retzius cells in the superficial layer of the developing forebrain, Reelin binds to the ApoER2 and VLDLR receptors on migrating neurons. Reelin binding to its receptors induces clustering of its downstream adaptor protein, Dab1, which is then phosphorylated by SRC family tyrosine kinases (Hiesberger et al., 1999; Benhayon et al., 2003). Phosphorylation of Dab1 promotes its adaptor function to recruit downstream factors that mediate neuronal migration and cell layering in the forebrain. Accordingly, loss of Dab1, ApoER2, or VLDLR function mimics the phenotypes of the *reeler* mutant (Trommsdorff et al., 1999; Sharaf et al., 2013). Altogether, many studies have established Reelin as an important cue for migrating neurons.

The Reelin/Dab1 signaling pathway has additionally been implicated in the maturation of neurons after migration. *reeler* and *Dab1* knockout mutant mice exhibit reduced elaboration of neuronal dendritic arbors and dendritic spine development, phenotypes that can be rescued by Reelin supplementation of neurons derived from *reeler* brains (Niu et al., 2004; Niu et al., 2008). Importantly, neurons in heterozygous *reeler* mice have reduced dendrite and dendritic spine development, despite correct neuronal positioning. Furthermore, conditional knockout of *Dab1* in correctly positioned forebrain neurons reduces their dendritic complexity and synaptic connectivity (Matsuki et al., 2008; Ventruti et al., 2011; Trotter et al., 2013; Bosch et al., 2016). Altogether, these findings indicate that Reelin influences significant scopes of neuronal maturation and function independent of its function in cell migration and layering. However, the molecular factors that act downstream of Reelin/Dab1 that mediate these aspects of neuronal maturation have yet to be fully explored.

Here, we identify a novel interaction between Dab1 and Csmd2, a synaptic transmembrane protein that we recently demonstrated plays a role in dendrite and dendritic spine development and stability (Gutierrez et al., 2019). We show that Dab1 binds the cytoplasmic tail of Csmd2, suggesting a potential role for Csmd2 in the Reelin signaling cascade. In determining whether Csmd2 requires the intracellular Dab1-interacting motif to mediate structural development, we first show that both a wild-type version of Csmd2 and a mutant that is unable to interact with Dab1 are both able to localize to postsynaptic dendritic spines of forebrain neurons at comparable levels. However, expression of the mutant form of Csmd2 that is unable to interact with Dab1 results in an overproduction of dendritic spines that are of an immature, filopodia-like morphology. This suggests that the Dab1-interacting motif of Csmd2 is required for the structural maturation of dendritic spines. Finally, we find that Csmd2 is required for the ability of Reelin to promote the development of dendrites and dendritic protrusions in dissociated hippocampal neuron cultures. Together, these data suggest a role for Csmd2 in the Reelin/Dab1 signaling pathway to promote elaboration of the dendritic arbor, development of dendritic spines, and structural maturation of synapses via an interaction with Dab1.

## Materials and Methods

### Animals

All experimental methods used in this study were approved by the Institutional Animal Care and use Committee of the University of Colorado Denver, Anschutz Medical Campus, and were performed according to approved protocols and guidelines. All experiments involving mouse tissue, unless otherwise indicated, were conducted using hybrid F1 mice resulting from crosses between *129X1/SvJ* (https://www.jax.org/strain/000691, RRID: MGI:5653118) and *C57BL/6J* (https://www.jax.org/strain/000664, RRID: MGI:5656552). Mice of either sex that resulted from the above crosses were used in this study.

### DNA Plasmid Constructs

Partial gene fragments of mouse *Csmd2* cDNA were amplified using reverse-transcription polymerase chain reaction from total RNA extracted from adult mouse cerebral cortex. Remaining fragments were synthesized and purchased from Integrated DNA Technologies (IDT) as gBlocks. Truncated 15x *Csmd2* cDNA was cloned using NEBuilder HiFi (New England Biolabs, E2621) into an expression vector comprising a CMV promoter and chicken beta-actin enhancer (CAG), the preprotrypsin (PPT) leader sequence and three tandem FLAG epitopes (3xFLAG). Each expression construct was cloned so that the PPT leader sequence and 3xFLAG were fused in frame at the N-terminus of Csmd2. Mutation of the C-terminal NPxY consensus motif was achieved by synthesizing the mutant cytoplasmic domain as a gBlock and cloning into the wild-type construct.

shRNA plasmids used in this study contained both an shRNA expression cassette and a reporter gene expression cassette. The shRNA sequences were synthesized as gBlocks (IDT) and cloned upstream of a U6 promoter. shRNA sequences were: Non-targeting control – 5’GCGATAGCGCTAATAATTT3’; *Csmd2* shRNA #1 – 5’ GGCAAAGTCCTCTACTGAA3’; *Csmd2* shRNA #2 – 5’GGACGTTCTTCAGATATAA3’. The reporter gene encoded a myristoylated form of TdTomato that targets the fluorescent reporter to the plasma membrane, which was cloned downstream of the CAG promoter/enhancer. Unless indicated, a combination of *Csmd2* shRNAs #1 and #2 were used for knockdown of *Csmd2* mRNA, which we’ve previously validated and utilized for greater specificity while minimizing off-target effects (Gutierrez et al., 2019).

All constructs were confirmed by DNA sequencing. Detailed methods and maps for all expression vectors will be provided upon request.

### mRNA In Situ Hybridization

E14.5 brains were cut coronally on a cryostat at 12 μm. mRNA *in situ* hybridizations for *Csdm1, Csmd2* and *Csmd3* were carried out as previously described (Franco et al., 2012). The *Dab1 in situ* hybridization image was taken from the GenePaint publically available database (https://gp3.mpg.de) (Visel et al., 2004) and is a sagittal section from an E14.5 embryo: set ID MH356, image C997.6.1.B. cDNAs used to make probes were cloned into pBluescript II SK+ using primers: Csmd1 forward 5’-GGGCATCCATCAACTACTA-3’, reverse 5’-TTGGGTGTGGCTGTGAGATAAG-3’; Csmd2 forward 5’-GGAAGGCACTCGAAGGTCACTG-3’, reverse 5’-AGAAGTGTTTGACGGAGCAGATAA-3’; Csmd3 forward 5’-CAGCAACAAAAACTGGCTCA-3’, reverse 5’-TCACTCCAAGCAGCAAACAC-3’.

### 3x FLAG Pull-Down and Co-Immunoprecipitation

For FLAG pull-down and co-immunoprecipitation experiments, samples were lysed in a working solution of 50 mM Tris-HCl, 1 mM NaCl, 1% Triton X-100, and 1 mM EDTA, pH 7.6. Every 10 mL of this working solution was supplemented with 1 cOmplete ULTRA, Mini, EDTA-free protease inhibitor cocktail tablet (Roche, 11836170001). Samples were incubated for 3 hours at 4°C with Anti-DYDDDDK Affinity Gel (Rockland, RRID:AB_10704031). After incubation, washes were conducted according to the manufacturer’s recommended protocol. Samples were eluted from beads via incubation with Laemmli sample buffer (Bio-Rad, 1610737) at 37°C for 20 minutes prior to western blot analysis.

### Western Blot Analysis

Protein Concentrations were measured with the BCA assay (Pierce, Thermo Scientific, 23225) prior to SDS-PAGE and western blot. All protein samples were subjected to SDS-PAGE using 4-15% polyacrylamide gradient gels (Bio-Rad, 4561086). For cell lysates, 20-30 μg was loaded, while for synaptosomal fraction samples 40 μg of protein for each sample was loaded on the gels. Separated proteins were then electroblotted using a TransBlot Turbo system to TransBlot Turbo Mini-size PVDF membranes (Bio-Rad, 1704272). Membranes were subsequently blocked with 1x TBS containing 0.1% Tween 20 (1x TBST) with 5% (w/v) blotting-grade blocker (Bio-Rad, 1706404) and probed with the primary antibody of interest diluted in 1x TBS containing 0.1% Tween 20 and 0.5% blocker at room temperature overnight. Primary antibodies used: Dab1 (Millipore, RRID: AB2261451) 1:1000, the DYKDDDK epitope (FLAG; ThermoFisher, RRID: AB_2536846) 1:500, and Csmd2 (Novus Biologicals, RRID: AB_11019509) 1:200. Membranes were washed 3 times in 1x TBST for 10 minutes each prior to 1 hour of incubation at room temperature with the appropriate horseradish peroxidase (HRP)-conjugated secondary antibodies used at 1:1000. Membranes were visualized using the Clarity Western ECL Blotting Substrates (Bio-Rad, 1705060) according to the manufacturer’s recommended instructions in a Bio-Rad Chemidoc Universal Hood III imaging system.

### In Utero Electroporation

*In utero* electroporations were performed as described (Franco et al., 2011). Briefly, timed pregnant mice (E14.5 or E15.5) were anesthetized and their uterine horns exposed. 1-2 μL of endotoxin-free plasmid DNA was injected into the embryos’ lateral ventricles at 2 mg/mL each. For electroporation, 5 pulses separated by 950 ms were applied at 50 V. Embryos were allowed to develop *in utero* and postnatally until the indicated timepoints.

### Immunohistochemistry

Mouse brains were fixed by transcardial perfusion with 4% paraformaldehyde before dissection and additional post-fixation for 3 hours at room temperature. Free-floating coronal sections were cut at 75 μm on a vibratome. Prior to immunohistochemical analyses, sections were subjected to antigen retrieval by incubation in 10 mM sodium citrate, pH 6.0 in a pressure cooker set to cook at pressure for 1 minute.

For immunohistochemistry, sections were rinsed with 1x PBS twice for 5 minutes each. Sections were permeabilized and blocked with 10% normal donkey serum (Jackson ImmunoResearch, RRID: AB_2337254) and 0.1% Triton X-100 (Sigma-Aldrich) in 1x PBS for 1 hour at room temperature. Primary antibodies used: Tbr1 (Millipore, RRID: AB_10806888) 1:500, Ctip2 (Abcam, RRID: AB_2064130) 1:1000, Cux1 (Santa Cruz Biotechnology, RRID: AB_2261231) 1:200, Csmd2 (Novus Biologicals, RRID: AB_11019509) 1:200, GFP (ThermoFisher, RRID: AB_221569) 1:500, and the DYKDDDK epitope (FLAG; ThermoFisher, RRID: AB_2536846) 1:200.

### Synaptosomal Fractionation

Preparation of crude synaptosomal fractions and crude postsynaptic density fractions were prepared from mouse forebrain homogenate was performed as previously described (Sanderson et al., 2012). Samples obtained from the fractions produced by this protocol were subjected to SDS-PAGE and western blotting as described above.

### Primary Hippocampal Neuron Culture

Primary cultured hippocampal neurons were prepared from embryonic day (E) 17.5 *C57Bl/6J* mice. Hippocampal tissue was manually dissected and dissociated as previously described (Lesuisse and Martin, 2002). 500,000 cells were seeded per well onto poly-D-lysine-coated (Millipore, A-003-E) 12 mm cover slips in 24-well plates in Dulbecco’s Modification of Eagle’s Medium (DMEM; Corning, 10-017-CV) containing 10% fetal bovine serum (Gibco, 10437010) and 1% penicillin/streptomycin (Lonza, 17-602E). At 1 day *in vitro* (DIV), the DMEM-based culture medium was replaced with Minimum Essential Eagle’s Medium (EMEM; Lonza, 12-125F) containing 2.38 mM sodium bicarbonate (Sigma, S5761-500G), 2 mM stabilized L-glutamine (Gemini Bio, 400-106), 0.4% glucose (Sigma, G7021-100G), 0.1 mg/mL apo-transferrin (Gemini Bio, 800-130P), 2% Gem21 NeuroPlex Serum-Free Supplement (Gemini Bio, 400-160), 5% fetal bovine serum (Gibco, A31604-01), and 1%penicillin/streptomycin (Lonza, 17-602E). Half of the culture medium was replaced with fresh complete EMEM-based medium every other day. At the indicated times, coverslips were fixed in 4% paraformaldehyde for 20 minutes at room temperature, washed 3x with PBS, and mounted onto microscope slides using ProLong Diamond antifade reagent (Invitrogen, P36970).

### Transfection of Primary Hippocampal Neuron Cultures

Upon dissection and dissociation of hippocampal tissue as described above, cells were transfected with the Amaxa Mouse Neuron Nucleofector Kit (Lonza, VPG-1001) using the Amaxa Nucleofector II device (Lonza). Transfection was conducted according to the manufacturer’s recommended protocol for primary mouse hippocampal and cortical neurons.

### Preparation of Reelin-containing media

Reelin-containing and control conditioned media were obtained by collection of conditioned media from the stable cell line CER and the HEK293-EBNA cell lines, respectively (Niu et al., 2004; Niu et al., 2008), which were graciously provided by Dr. Gabriella D’Arcangelo and her research group in the Department of Cell Biology and Neuroscience at Rutgers University. Both cell lines were maintained in Dulbecco’s Modification of Eagle’s Medium (DMEM; Corning, 10-017-CV) containing 10% fetal bovine serum (Gibco, 10437010), 2 mM stabilized L-glutamine (Gemini Bio, 400-106), and 1% penicillin/streptomycin (Lonza, 17-602E). Cells were transitioned to serum-free DMEM-based media for collection of conditioned media from both lines. Conditioned media from both cell lines were collected every 3 days for 6 days. Collected media was centrifuged at 1500 RPM for 5 minutes to pellet dead cells. Supernatant was collected and used in 50% dilution with EMEM-based hippocampal neuron media for Reelin treatments. Recombinant Reelin was purchased from R&D Systems (R&D Systems, 3820-MR), reconstituted in EMEM and used at the indicated concentration in EMEM-based hippocampal neuron media.

### Statistical Analysis

Dendritic spine densities and morphological analyses were performed using ImageJ. All quantitative data were graphed as the mean with the standard error of the mean (SEM) of each experimental group. Student’s t-test was used to consider statistical significance, which was determined when p<0.05.

## Results

### Csmd2 interacts with Dab1 via FENPMY motif on intracellular Csmd2

The N-terminus of Dab1 contains a domain that can simultaneously bind membrane phosphoinositides and NPxY peptide motifs (Fig. 1A), allowing Dab1 to bind with high affinity to membrane-anchored proteins containing this motif in their cytoplasmic tails (Howell et al., 1999; Stolt et al., 2003; Stolt et al., 2004; Stolt et al., 2005). This mechanism of plasma membrane targeting is reported to be crucial for Reelin-mediated phosphorylation of Dab1 and its subsequent interaction with components of the Reelin cascade (Howell et al., 1999). Structural studies and protein interaction data have identified F/Y-D/E-NPxY as the Dab1 consensus binding motif (Howell et al., 1999; Stolt et al., 2003). We reasoned that other proteins containing this sequence might be novel Dab1 interactors, so we searched the NCBI RefSeq and EMBL-EBI protein databases for proteins containing the motif. In addition to several known Dab1-interacting proteins, we found that Csmd proteins contain consensus Dab1-binding motifs in their cytoplasmic tails (Fig. 1B). Therefore, we hypothesized that the Csmd family of proteins may be downstream components of the Reelin/Dab1 signaling cascade.

**Figure 1.**
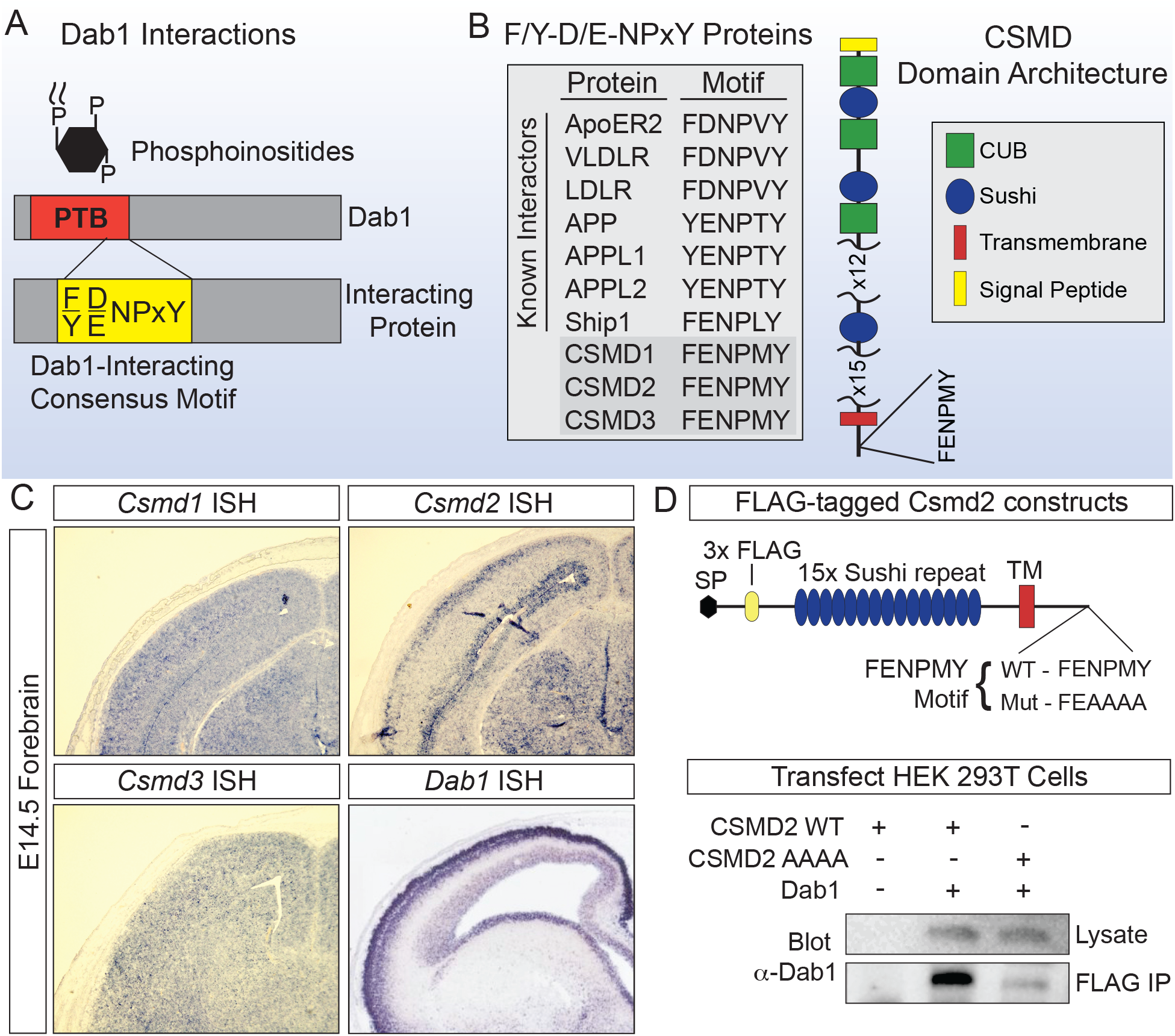
Dab1 interacts with Csmd2 via FENPMY motif. **A.** Schematic of Dab1 protein, noting the phosphotyrosine binding (PTB) domain through which Dab1 docks to the plasma membrane and simultaneously binds F/Y-D/E-NPxY motifs of protein interactors. **B.** Known interactors of Dab1 and their corresponding Dab1-binding motifs. Csmd protein family members each contain an FENPMY motif. **C.** mRNA *in situ* hybridization of E14.5 mouse forebrains comparing expression of *Csmd1*, *Csmd2*, and *Csmd3* to that of *Dab1*. Of the three *Csmd* mRNAs, the *Csmd2* mRNA expression pattern appears to match that of *Dab1* mRNA most closely. *Dab1 in situ* image taken from the GenePaint database, set ID MH356, image C997.6.1.B. **D.** Schematic of FLAG-tagged Csmd2 constructs generated for *in vitro* characterization of Csmd2-Dab1 interaction. Immunoprecipitation of FLAG-Csmd2 from transfected HEK293T cells pulls down co-transfected Dab1, but this interaction is abolished when the NPMY motif in Csmd2 is mutated to AAAA.

Among the three members of the Csmd gene family, we found that *Csmd2* mRNA was expressed in a pattern similar to that of *Dab1* in the developing forebrain (Fig. 1C). For this reason, we focused our attention on the relationship between Dab1 and Csmd2 in this study. To test whether Dab1 can interact with the FENPMY motif on the Csmd2 cytoplasmic tail, we cloned an expression construct comprising the Csmd2 cytoplasmic tail, transmembrane domain and a truncated extracellular domain containing the first 15 Sushi repeats and an N-terminal FLAG tag (hereafter referred to as Csmd2-15x). In parallel, we cloned a version of Csmd2-15x in which we mutated the FENPMY motif to FEAAAA (Fig. 1D). We then separately transfected each of these constructs, together with an expression vector for Dab1, into HEK293T cells (Fig. 1D). Upon co-immunoprecipitation of the 3xFLAG epitope on Csmd2-15x and subsequent Western blot for Dab1, we found that Dab1 was coimmunoprecipitated with wild-type Csmd2-15x, but not with the Csmd2-15x NPxY mutant (Fig. 1D). Therefore, we concluded that Dab1 can bind the Csmd2 cytoplasmic tail through the FENPMY motif.

### The Csmd2-Dab1 Interaction is Required for the Structural Maturation of Dendritic Spines

We previously showed that Csmd2 interacts with PSD95 through a PDZ-binding domain on the Csmd2 cytoplasmic tail, and that this interaction is required for Csdm2 to localize to synapses (Gutierrez et al., 2019). We wondered whether the interaction between Dab1 and Csmd2 would be similarly required for targeting Csmd2 to synapses. To address this question, we utilized the Csmd2-15x wild-type and NPxY mutant constructs (Fig. 1C, 2A). We introduced these constructs separately into mouse forebrain progenitors at E14.5 by *in utero* electroporation and then analyzed their localizations by biochemical fractionation and immunohistochemistry in immature (P7) and matured (P60) forebrains (Fig. 2A). We isolated postsynaptic density (PSD) fractions from crude synaptosome preparations and probed for the FLAG-tagged Csmd2-15x proteins by Western blot (Fig. 2B). We found that both the Csmd2-15x wild-type and NPxY mutant proteins localized to PSD fractions with no significant difference observed between them at either timepoint (Fig. 2B). These findings were further verified by immunohistochemical localization of the Csmd2-15x constucts in electroporated brains. We found that the Csmd2-15x NPxY mutant exhibited similar localization at PSD-95+ dendritic spines as the Csmd2-15x wild-type protein (Fig. 2C). Therefore, we concluded that Dab1 binding to the intracellular region of Csmd2 is not crucial for the localization of Csmd2 at synapses.

**Figure 2.**
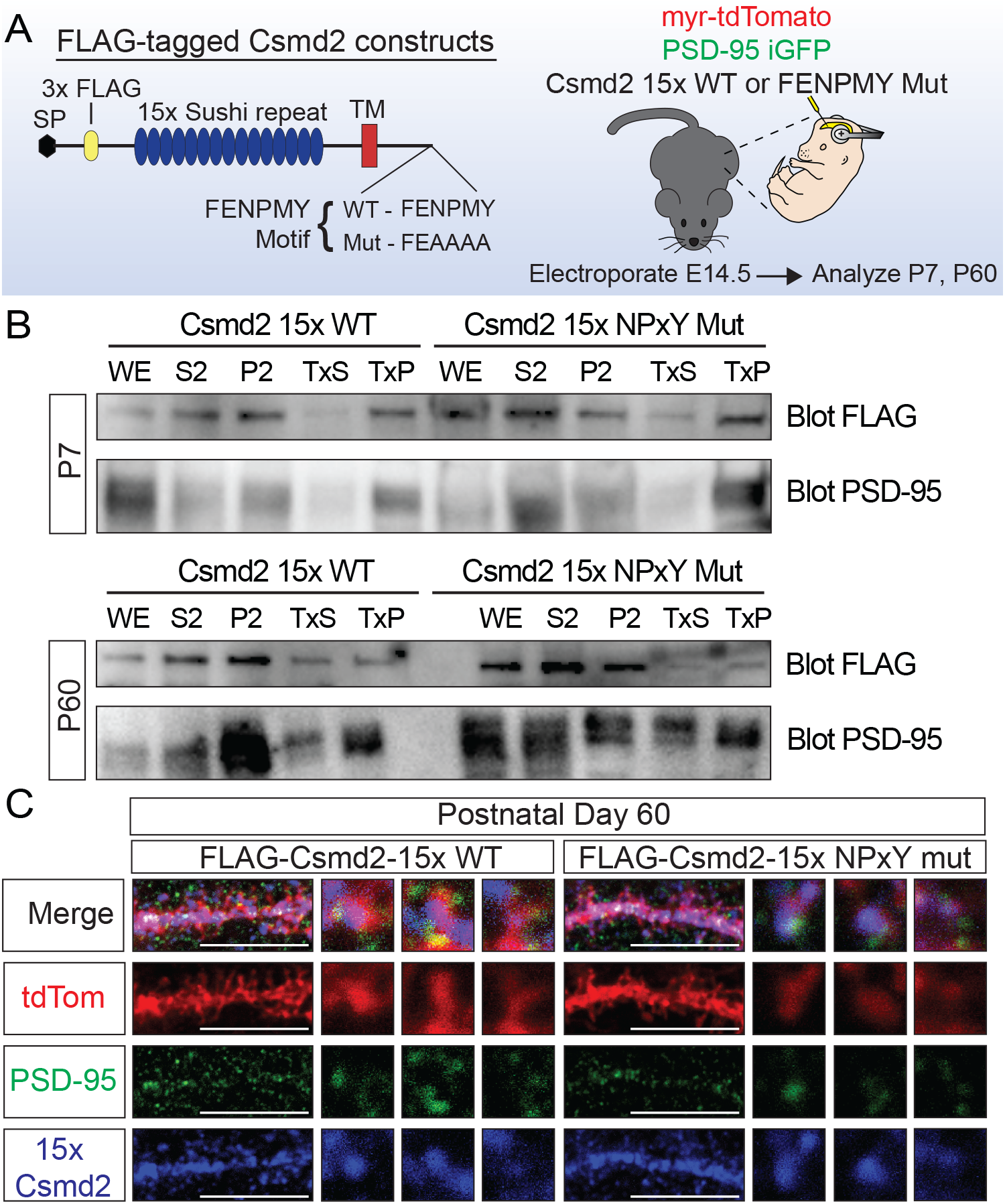
Mutation of Dab1-binding region does not affect the localization of Csmd2 at postsynaptic dendritic spines. **A.** Experimental design to visualize the localization of Csmd2 upon mutation of Dab1-binding FENPMY motif. FLAG-tagged wild-type and FENPMY mutant Csmd2 constructs were introduced into mouse forebrain progenitors at E14.5 by *in utero* electroporation. Electroporated brains were subsequently used at P7 or P60 for biochemical and immunohistochemical analyses. **B.** PSD-fraction enrichment by Triton X-100 extraction of crude synaptosomal fractions obtained from experiments outlined in **A.** Western blots reveal no difference in PSD enrichment between FLAG-Csmd2 wildtype and FENPMY mutant constructs. C. Immunohistochemical analysis for Csmd2 and PSD-95 confirmed no difference in dendritic spine localization between the two experiment groups. Scale bars = 10 μm.

While analyzing these electroporation experiments, we noticed that the dendritic protrusions on cells expressing the Csmd2-15x NPxY mutant appeared different from those on cells expressing the wild-type Csmd2-15x construct (Fig. 3A,B). Quantification of dendritic spines in each case showed that neurons expressing the Csmd2-15x NPxY mutant construct displayed a more than 2-fold increase in relative dendritic spine density compared to neurons expressing the wild-type Csmd2-15x construct (Fig. 3C). Interestingly, many of these increased dendritic spines were of a predominantly filopodia-like morphology, reminiscent of immature dendritic spines compared to more robust mushroom-shaped spines. Dendritic spines on Csmd2-15x NPxY mutant neurons displayed a 20% decrease in average spine head width (Fig. 3D). Average spine length was also modestly increased in Csmd2-15x NPxY mutant dendritic spines (Fig. 3E), consistent with less mature dendritic protrustions. These data raised the interesting possbility that Dab1 binding to the Csmd2 NPxY motif may be important for regulating the maturation of neuronal dendrites and synapses.

**Figure 3.**
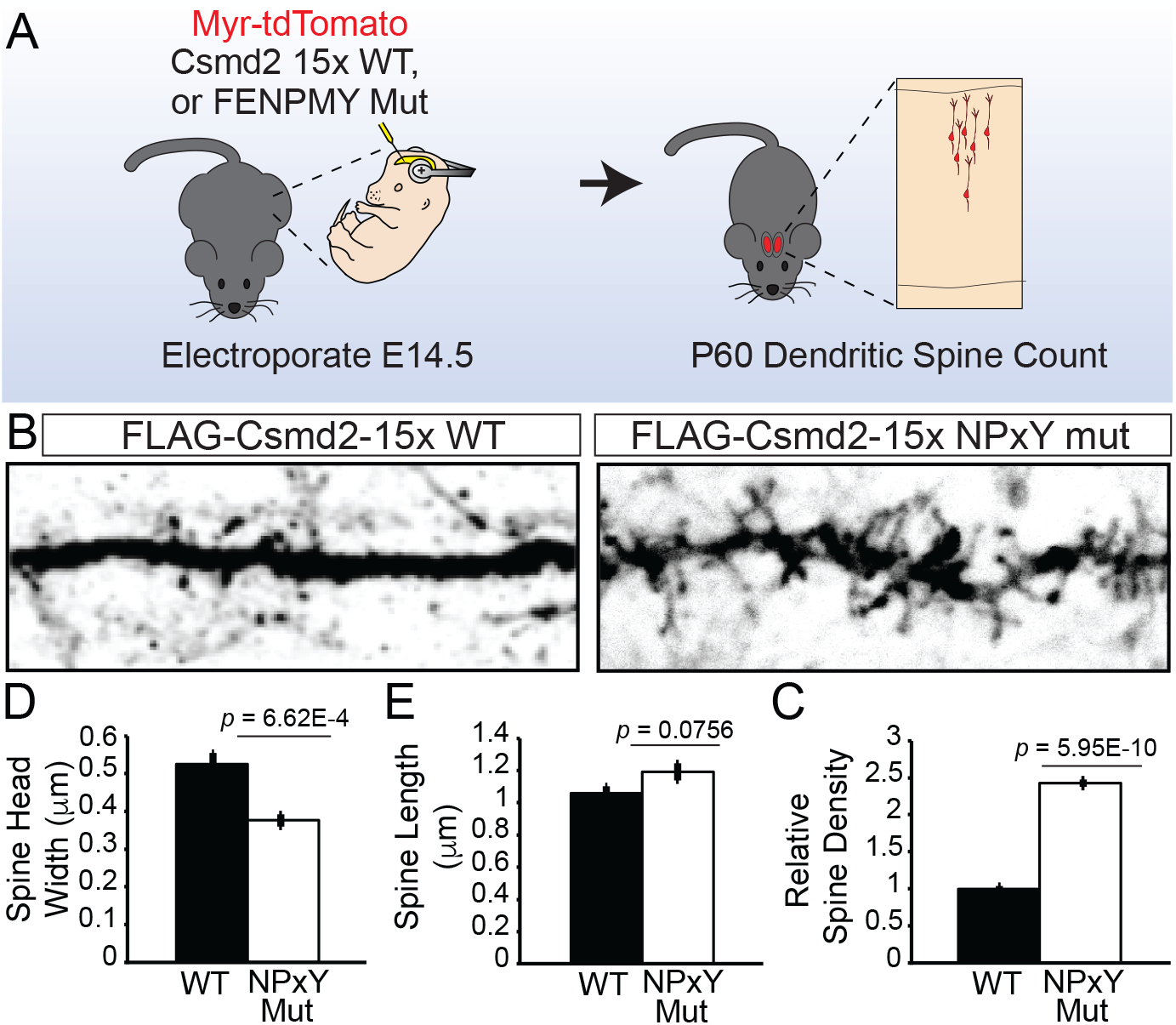
Introduction of Csmd2 NPxY mutant results in a prevalence of filopodia-like dendritic spines. **A.** Experimental design to determine the effect of Csmd2 NPxY mutant expression on dendritic spine density and structure in matured forebrain neurons. **B**. Representative images at P60 of dendrites from neurons expressing the versions of the truncated Csmd2 protein, as indicated. Compared to Csmd2-15x WT-expressing neurons, neurons expressing the Csmd2-15x NPxY mutant display increased dendritic spine densities. **C.** Quantification of dendritic spine density from (B). **D-E.** Measurements of dendritic spine head width (D) and spine length (E) show a prevalence of thinner spines upon expression of the Csmd2-15x NPxY mutant construct compared to that of the Csmd2-15x WT construct.

### Csmd2 is required for Reelin-mediated neuronal structural development

The Reelin signaling pathway plays a significant role in the elaboration of the neuronal dendritic arbor and development of postsynaptic dendritic spines (Niu et al., 2004; Jossin and Goffinet, 2007; Niu et al., 2008; Chai et al., 2015; Kim et al., 2015). Supplementation of Reelin has also been shown to enhance the structural development of neurons in these contexts (Niu et al., 2004; Niu et al., 2008; Rogers et al., 2011; Rogers et al., 2013), resulting in enhanced dendritic branching and outgrowth, increased dendritic spine density, and increased long-term potentiation. We recently showed that Csmd2 plays a role in dendrite and dendritic spine development and stability (Gutierrez et al., 2019). Given the interaction between Dab1 and Csmd2, we asked whether Csmd2 is required downstream of Reelin to mediate the development of dendritic spines and elaboration of the dendritic arbor. To address this question, we supplemented primary hippocampal neuron cultures with either control (CTRL) or Reelin-containing conditioned media and assessed dendrite development in the context of either shRNA-mediated *Csmd2* knockdown (shCsmd2) or non-targeting control shRNA (shCTRL) conditions. After 5 days of conditioned media treatment starting at the day of plating (0 days *in vitro*, 0 DIV), neurons cultured with Reelin-containing conditioned media displayed approximately 30% more dendritic protrusions per micrometer of dendrite compared to neurons cultured with CTRL conditioned media (Fig. 4), in line with previous reports (Niu et al., 2008; Rogers et al., 2011; Hethorn et al., 2015). However, upon knockdown of *Csmd2* mRNA expression, this effect was completely negated. *Csmd2* knockdown resulted in significantly fewer dendritic protrusions in control media treatment groups, as well as a failure of Reelin to increase the number of dendritic protrusions upon treatment with Reelin-containing conditioned media (Fig. 4). This effect was observed using two different *Csmd2*-targeting shRNAs (shCsmd2 #1, shCsmd2 #2), as well as when we combined both shRNAs each at half amounts (shCsmd2 #1+2) to control for potential off-target effects (Fig. 4).

**Figure 4.**
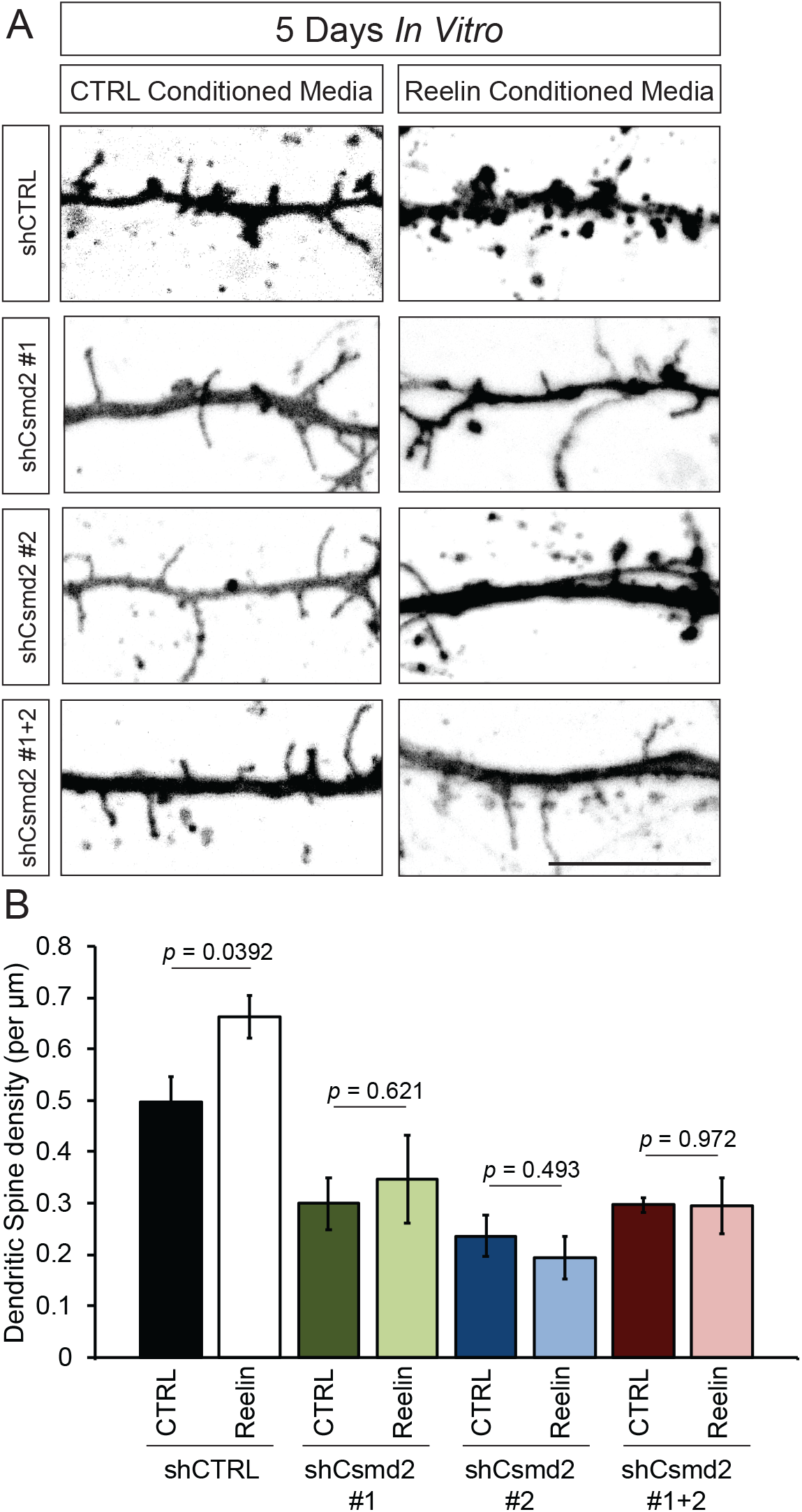
Knockdown of *Csmd2* mRNA disrupts Reelin-mediated dendritic spine development in cultured hippocampal neurons. **A.** Representative dendrite images from cultured mouse hippocampal neurons transfected upon plating (DIV 0) with *Csmd2*-targeting shRNAs together with treatment with conditioned media from either a control or Reelin-secreting line (0-5 DIV; scale bars = 10 μm). **B.** Quantification of dendritic filopodia per μm of dendrite. Treatment with Reelin-containing conditioned media increases the density of dendritic filopodia in control neurons expressing non-targeting shRNA, but not in neurons in which *Csmd2* mRNA is knocked down.

We next evaluated the requirement of Csmd2 in Reelin-mediated development of dendrite branching. After 5 days of conditioned media treatment starting at 0 DIV, we evaluated early dendrite complexity between shCTRL and shCsmd2 experiment groups by Sholl analysis. In line with previous reports (Niu et al., 2004), treatment of cultured neurons with Reelin-containing conditioned media resulted in an increase in early dendrite complexity compared to neurons treated with CTRL conditioned media (Fig. 5A,B). Conversely, Reelin-containing conditioned media treatment had no effect on early dendrite complexity in neurons in which *Csmd2* mRNA expression was knocked down (Fig. 5A,C). Even after chronic treatment with recombinant Reelin for 12 DIV,neurons in which *Csmd2* was knocked down failed to increase their dendrite complexity in response to Reelin treatment (Fig. 5D). Altogether, we concluded that *Csmd2* expression is critical for early dendrite elaboration and dendritic filopodia development promoted by Reelin supplementation.

**Figure 5.**
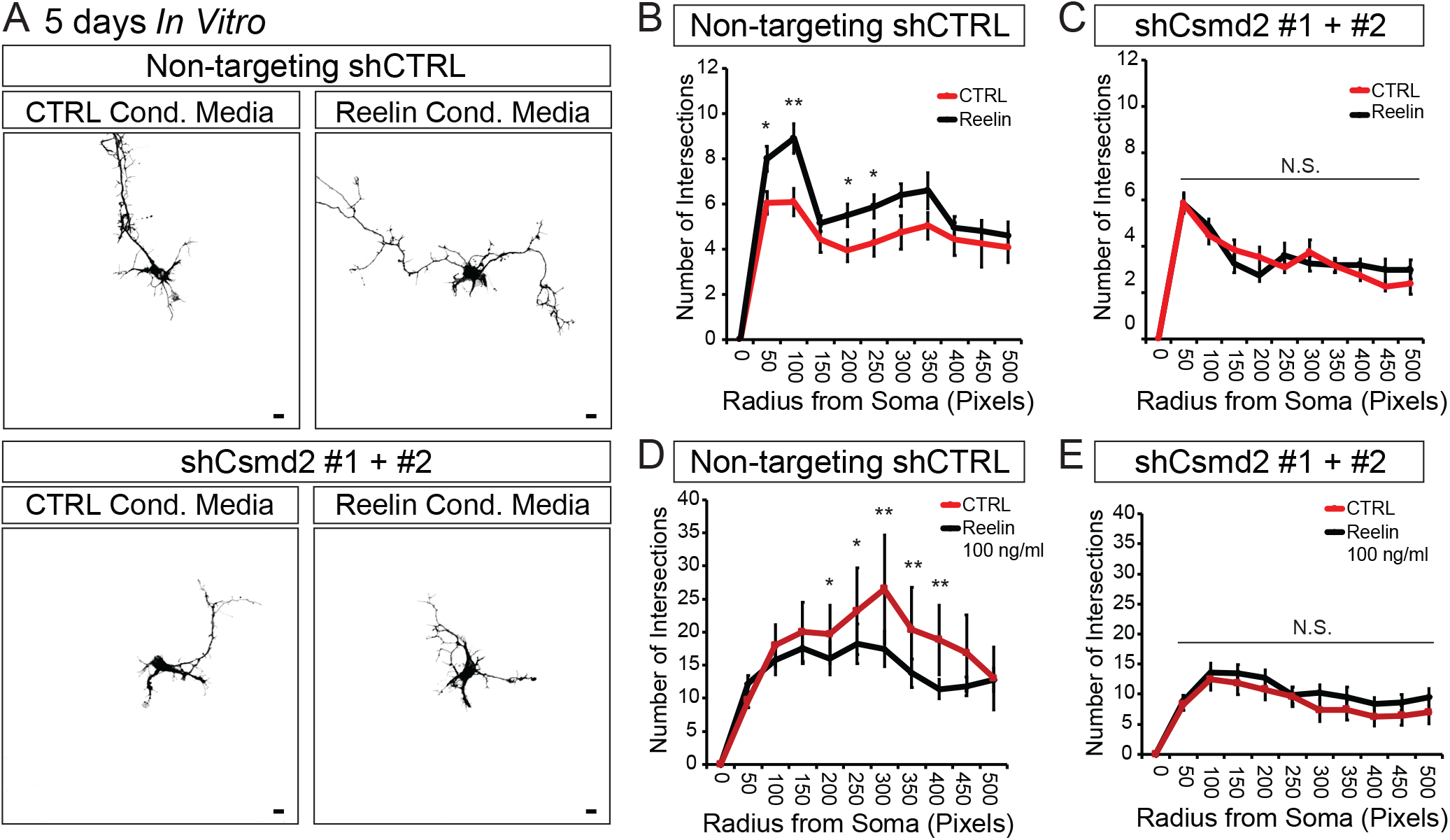
Knockdown of *Csmd2* mRNA disrupts Reelin-mediated dendrite development in cultured hippocampal neurons. **A.** Representative images of cultured mouse hippocampal neurons transfected upon plating (0 DIV) for the expression of *Csmd2*-targeting shRNAs and treated with conditioned media from either a control or Reelin-secreting cell line (0-5 DIV; scale bars = 10 μm). **B.** Quantification of dendrite complexity by Sholl analysis shows increased elaboration of neuronal dendritic arbors upon Reelin supplementation in control neurons. This effect was disrupted upon *Csmd2* mRNA knockdown, despite Reelin supplementation (C). **D.** Long-term treatment of cultured hippocampal neurons with recombinant Reelin (0-12 DIV; 100 ng/mL) resulted in increased dendrite complexity, as shown by Sholl analysis at 12 DIV. **E.** This increase in dendrite complexity after Reelin treatment was disrupted upon *Csmd2* mRNA knockdown.

## Discussion

Reelin loss-of-function mutations result in deficits in brain development, including in the structural development and maturation of neuronal dendrites and synapses. Such deficits are associated with neuropsychiatric disorders that exhibit cognitive and social setbacks such as schizophrenia and autism spectrum disorder, which can greatly affect an individual’s quality of life.

Here, we identify an interaction between the synaptic transmembrane protein, Csmd2, and the Reelin downstream adapter protein Dab1. Csmd2 contains an F/ENPxY consensus Dab1-binding motif in its cytoplasmic tail, which we find is required for the Dab1-Csmd2 interaction. We previously showed that the Csmd2 cytoplasmic tail also contains a consensus PDZ-binding motif that mediates interaction with the synaptic scaffold protein, PSD-95, and is important for synaptic localization of Csmd2 (Gutierrez et al., 2019). Unlike the PDZ-binding domain, we find that the NPxY domain of Csmd2 is not necessary for targeting Csmd2 to synapses. Although these data suggest that the interaction between Dab1 and Csmd2 is not required generally for Csdm2 localization at synapses, further studies will be needed to determine if more subtle changes in Csmd2 trafficking or subcellular localization are affected by its interaction with Dab1. This will be particularly interesting in the context of activated Reelin signaling, since Reelin is known to promote interactions between Dab1 and its transmembrane binding partners to regulate their cell surface localization (Hoe et al., 2006).

Given our previous report that Csmd2 is required for dendritic spine development and maintenance as well as elaboration of the dendritic arbor, we hypothesized that Csmd2 is a critical factor required downstream of Reelin/Dab1 signal transduction to mediate the structural maturation of a neuron. In line with this, we found that knockdown of *Csmd2* mRNA expression resulted in a failure of Reelin to enhance the early development of dendrites and dendritic filopodia. We further show that neurons expressing a mutated form of Csmd2 that no longer interacts with Dab1 present an overproduction of filopodia-like dendritic spines compared to neurons expressing the wild-type form. Taken together, these data suggest a model in which the Dab1-Csmd2 interaction mediates a downstream mechanism that governs dendritic spine development and maturation. While the Reelin/Dab1 signaling cascade plays a critical role in neuronal migration and layering during brain development, shRNA-mediated knockdown of *Csmd2* mRNA in immature migrating neurons exhibit no defects in neuronal migration and placement (unpublished data, M.A. Gutierrez & S.J. Franco). This suggests a role for Csmd2 as a molecular factor that governs a function of the Reelin/Dab1 cascade separately from neuronal migration.

The molecular mechanism through which the Dab1-Csmd2 interaction mediates synapse development and maturation remain to be elucidated. We may gain some insight through previous reports describing how Reelin regulates the localization of ionotropic glutamate receptors and its influence on synaptic activity and long-term potentiation (LTP). One interesting avenue of study in this regard is the potentiation of calcium influx by Reelin through NMDA receptors (NMDARs). Reelin has been previously reported to induce tyrosine phosphorylation of the cytoplasmic NMDAR subunit NR2B through SRC family kinases, thus increasing ion conductance at the synapse and ultimately LTP (Chen et al., 2005). This enhanced ion conductance of synaptic NMDARs results in greater calcium influx which, in turn, results in enhanced dendritic spine density and development through Calcium/calmodulin-dependent protein kinase type II β (CaMKIIβ) kinase activity and transcriptional activity of cAMP-response element binding protein (CREBP) (Chen et al., 2005; Kim et al., 2015). It would be interesting to address whether Csmd2 is required for this plasticity in synaptic physiology induced by Reelin; in particular, whether the Csmd2-Dab1 interaction mediates downstream CaMKIIβ and CREBP signaling.

Other CUB/Sushi-containing proteins have been described in previous reports as auxiliary subunits of synaptic ionotropic glutamate receptors that regulate receptor subunit composition, localization, clustering, and transport throughout the dendritic arbor and finally to the synapse (Gendrel et al., 2009; Ng et al., 2009; Cousins et al., 2013; Fisher and Mott, 2013; Lomash et al., 2017; Wyeth et al., 2017). Additionally, Reelin has been reported to induce signal transduction through phosphoinositide-3 kinase (PI3K) to enhance surface insertion of AMPA receptors (AMPARs), contributing to an enhanced NMDAR response (Qiu et al., 2006). Therefore, it would be interesting to address whether Csmd2 is required in the transport, clustering, subunit composition, and synaptic mobility that would then influence ion conductance of NMDARs, AMPARs, and kainate receptors, and whether Csmd2 is required for the enhancement of these parameters by Reelin supplementation.

In this study, we focused our attention on the role of Csmd2 in the Reelin-mediated structural maturation of immature, developing neurons. However, it is also known that Reelin is expressed by GABAergic in terneuronsin the adult forebrain (Alcantara et al., 1998; Drakew et al., 1998; Pesold et al., 1998). Additionally, overexpression of Reelin in adult mouse forebrains results in the hypertrophy of dendritic spines and enhanced LTP (Pujadas et al., 2010). Therefore, it will be interesting to investigate whether Csmd2 is required for this enhancement of synaptic maturity in an adult system *in vivo*. In particular, it would be of significant clinical relevance to pursue this question in the context of a neurodegenerative, aging, or traumatic injury contexts in which synaptic stability is at risk. Our previous work indicates that Csmd2 loss of function in matured neurons *in vitro* results in significantly reduced dendritic spine density and dendritic complexity (Gutierrez et al., 2019). Therefore, it is possible that Csmd2 may indeed be a target in the Reelin/Dab1 pathway for the development of therapeutic strategies that can maintain synapse maturity and function in the adult brain.

In conclusion, we have identified an interaction between the synaptic transmembrane protein Csmd2 and the Reelin-associated adapter protein Dab1, as well as a requirement for Csmd2 in Reelin-mediated early dendritic elaboration and dendritic spine maturation. Future studies focusing on the requirement of Csmd2 in Reelin/Dab1 signal transduction and synaptic physiology will lead to an expansion in our understanding of how Reelin governs forebrain developmental dynamics and postnatal function. As such, these studies will help us better understand the molecular and cellular basis of neuropsychiatric disorders associated with cognitive and social deficits that can greatly affect an individual’s quality of life.

## Acknowledgements

We thank Drs. Mark Dell’Acqua, Matthew Kennedy, and Jason Aoto for reagents and technical advice. We also thank Dr. Gabriella D’Arcangelo for providing the Reelin-secreting cell line and technical advice described in the Materials and Methods section of this manuscript.

## Notes

**Conflict of Interest:** Authors report no conflict of interest.

**Funding sources:** This work was supported by NIH/NCATS Colorado CTSA Grant Number UL1 TR002535 (SJF), Children’s Hospital Colorado Program in Pediatric Stem Cell Biology (SJF) and The Boettcher Foundation (SJF).

## References

Alcantara S, Ruiz M, D’Arcangelo G, Ezan F, de Lecea L, Curran T, Sotelo C, Soriano E (1998) Regional and cellular patterns of reelin mRNA expression in the forebrain of the developing and adult mouse. J Neurosci 18:7779–7799.

Benhayon D, Magdaleno S, Curran T (2003) Binding of purified Reelin to ApoER2 and VLDLR mediates tyrosine phosphorylation of Disabled-1. Brain Res Mol Brain Res 112:33–45.

Bosch C, Masachs N, Exposito-Alonso D, Martinez A, Teixeira CM, Fernaud I, Pujadas L, Ulloa F, Comella JX, DeFelipe J, Merchan-Perez A, Soriano E (2016) Reelin Regulates the Maturation of Dendritic Spines, Synaptogenesis and Glial Ensheathment of Newborn Granule Cells. Cereb Cortex 26:4282–4298.

Chai X, Fan L, Shao H, Lu X, Zhang W, Li J, Wang J, Chen S, Frotscher M, Zhao S (2015) Reelin Induces Branching of Neurons and Radial Glial Cells during Corticogenesis. Cereb Cortex 25:3640–3653.

Chen Y, Beffert U, Ertunc M, Tang TS, Kavalali ET, Bezprozvanny I, Herz J (2005) Reelin modulates NMDA receptor activity in cortical neurons. J Neurosci 25:8209–8216.

Cousins SL, Innocent N, Stephenson FA (2013) Neto1 associates with the NMDA receptor/amyloid precursor protein complex. J Neurochem 126:554–564.

Drakew A, Frotscher M, Deller T, Ogawa M, Heimrich B (1998) Developmental distribution of a reeler gene-related antigen in the rat hippocampal formation visualized by Cr-50 immunocytochemistry. Neuroscience 82:1079–1086.

Eastwood SL, Harrison PJ (2006) Cellular basis of reduced cortical reelin expression in schizophrenia. Am J Psychiatry 163:540–542.

Falconer DS (1951) Two new mutants, ‘trembler’ and ‘reeler’, with neurological actions in the house mouse (Mus musculus L.). J Genet 50:192–201.

Fisher JL, Mott DD (2013) Modulation of homomeric and heteromeric kainate receptors by the auxiliary subunit Neto1. J Physiol 591:4711–4724.

Franco SJ, Martinez-Garay I, Gil-Sanz C, Harkins-Perry SR, Muller U (2011) Reelin regulates cadherin function via Dab1/Rap1 to control neuronal migration and lamination in the neocortex. Neuron 69:482–497.

Franco SJ, Gil-Sanz C, Martinez-Garay I, Espinosa A, Harkins-Perry SR, Ramos C, Muller U (2012) Fate-restricted neural progenitors in the mammalian cerebral cortex. Science 337:746–749.

Gendrel M, Rapti G, Richmond JE, Bessereau JL (2009) A secreted complement-control-related protein ensures acetylcholine receptor clustering. Nature 461:992–996.

Gutierrez MA, Dwyer BE, Franco SJ (2019) Csmd2 Is a Synaptic Transmembrane Protein that Interacts with PSD-95 and Is Required for Neuronal Maturation. eNeuro 6.

Hethorn WR, Ciarlone SL, Filonova I, Rogers JT, Aguirre D, Ramirez RA, Grieco JC, Peters MM, Gulick D, Anderson AE, J LB, Lussier AL, Weeber EJ (2015) Reelin supplementation recovers synaptic plasticity and cognitive deficits in a mouse model for Angelman syndrome. Eur J Neurosci 41:1372–1380.

Hiesberger T, Trommsdorff M, Howell BW, Goffinet A, Mumby MC, Cooper JA, Herz J (1999) Direct binding of Reelin to VLDL receptor and ApoE receptor 2 induces tyrosine phosphorylation of disabled-1 and modulates tau phosphorylation. Neuron 24:481–489.

Howell BW, Lanier LM, Frank R, Gertler FB, Cooper JA (1999) The disabled 1 phosphotyrosine-binding domain binds to the internalization signals of transmembrane glycoproteins and to phospholipids. Mol Cell Biol 19:5179–5188.

Jossin Y, Goffinet AM (2007) Reelin signals through phosphatidylinositol 3-kinase and Akt to control cortical development and through mTor to regulate dendritic growth. Mol Cell Biol 27:7113–7124.

Kim M, Jeong Y, Chang YC (2015) Extracellular matrix protein reelin regulate dendritic spine density through CaMKIIbeta. Neurosci Lett 599:97–101.

Lammert DB, Middleton FA, Pan J, Olson EC, Howell BW (2017) The de novo autism spectrum disorder RELN R2290C mutation reduces Reelin secretion and increases protein disulfide isomerase expression. J Neurochem 142:89–102.

Lesuisse C, Martin LJ (2002) Long-term culture of mouse cortical neurons as a model for neuronal development, aging, and death. J Neurobiol 51:9–23.

Lomash RM, Sheng N, Li Y, Nicoll RA, Roche KW (2017) Phosphorylation of the kainate receptor (KAR) auxiliary subunit Neto2 at serine 409 regulates synaptic targeting of the KAR subunit GluK1. J Biol Chem 292:15369–15377.

Matsuki T, Pramatarova A, Howell BW (2008) Reduction of Crk and CrkL expression blocks reelin-induced dendritogenesis. J Cell Sci 121:1869–1875.

Ng D, Pitcher GM, Szilard RK, Sertie A, Kanisek M, Clapcote SJ, Lipina T, Kalia LV, Joo D, McKerlie C, Cortez M, Roder JC, Salter MW, McInnes RR (2009) Neto1 is a novel CUB-domain NMDA receptor-interacting protein required for synaptic plasticity and learning. PLoS Biol 7:e41.

Niu S, Yabut O, D’Arcangelo G (2008) The Reelin signaling pathway promotes dendritic spine development in hippocampal neurons. J Neurosci 28:10339–10348.

Niu S, Renfro A, Quattrocchi CC, Sheldon M, D’Arcangelo G (2004) Reelin promotes hippocampal dendrite development through the VLDLR/ApoER2-Dab1 pathway. Neuron 41:71–84.

Olson EC, Kim S, Walsh CA (2006) Impaired neuronal positioning and dendritogenesis in the neocortex after cell-autonomous Dab1 suppression. J Neurosci 26:1767–1775.

Ovadia G, Shifman S (2011) The genetic variation of RELN expression in schizophrenia and bipolar disorder. PLoS One 6:e19955.

Pesold C, Impagnatiello F, Pisu MG, Uzunov DP, Costa E, Guidotti A, Caruncho HJ (1998) Reelin is preferentially expressed in neurons synthesizing gamma-aminobutyric acid in cortex and hippocampus of adult rats. Proc Natl Acad Sci U S A 95:3221–3226.

Pujadas L, Gruart A, Bosch C, Delgado L, Teixeira CM, Rossi D, de Lecea L, Martinez A, Delgado-Garcia JM, Soriano E (2010) Reelin regulates postnatal neurogenesis and enhances spine hypertrophy and longterm potentiation. J Neurosci 30:4636–4649.

Qiu S, Zhao LF, Korwek KM, Weeber EJ (2006) Differential reelin-induced enhancement of NMDA and AMPA receptor activity in the adult hippocampus. J Neurosci 26:12943–12955.

Rice DS, Sheldon M, D’Arcangelo G, Nakajima K, Goldowitz D, Curran T (1998) Disabled-1 acts downstream of Reelin in a signaling pathway that controls laminar organization in the mammalian brain. Development 125:3719–3729.

Rogers JT, Rusiana I, Trotter J, Zhao L, Donaldson E, Pak DT, Babus LW, Peters M, Banko JL, Chavis P, Rebeck GW, Hoe HS, Weeber EJ (2011) Reelin supplementation enhances cognitive ability, synaptic plasticity, and dendritic spine density. Learn Mem 18:558–564.

Rogers JT, Zhao L, Trotter JH, Rusiana I, Peters MM, Li Q, Donaldson E, Banko JL, Keenoy KE, Rebeck GW, Hoe HS, D’sArcangelo G, Weeber EJ (2013) Reelin supplementation recovers sensorimotor gating, synaptic plasticity and associative learning deficits in the heterozygous reeler mouse. J Psychopharmacol 27:386–395.

Sanchez-Sanchez SM, Magdalon J, Griesi-Oliveira K, Yamamoto GL, Santacruz-Perez C, Fogo M, Passos-Bueno MR, Sertie AL (2018) Rare RELN variants affect Reelin-DAB1 signal transduction in autism spectrum disorder. Hum Mutat 39:1372–1383.

Sanderson JL, Gorski JA, Gibson ES, Lam P, Freund RK, Chick WS, Dell’Acqua ML (2012) AKAP150-anchored calcineurin regulates synaptic plasticity by limiting synaptic incorporation of Ca2+-permeable AMPA receptors. J Neurosci 32:15036–15052.

Sharaf A, Bock HH, Spittau B, Bouche E, Krieglstein K (2013) ApoER2 and VLDLr are required for mediating reelin signalling pathway for normal migration and positioning of mesencephalic dopaminergic neurons. PLoS One 8:e71091.

Sobue A, Kushima I, Nagai T, Shan W, Kohno T, Aleksic B, Aoyama Y, Mori D, Arioka Y, Kawano N, Yamamoto M, Hattori M, Nabeshima T, Yamada K, Ozaki N (2018) Genetic and animal model analyses reveal the pathogenic role of a novel deletion of RELN in schizophrenia. Sci Rep 8:13046.

Stolt PC, Vardar D, Blacklow SC (2004) The dual-function disabled-1 PTB domain exhibits site independence in binding phosphoinositide and peptide ligands. Biochemistry 43:10979–10987.

Stolt PC, Jeon H, Song HK, Herz J, Eck MJ, Blacklow SC (2003) Origins of peptide selectivity and phosphoinositide binding revealed by structures of disabled-1 PTB domain complexes. Structure 11:569–579.

Stolt PC, Chen Y, Liu P, Bock HH, Blacklow SC, Herz J (2005) Phosphoinositide binding by the disabled-1 PTB domain is necessary for membrane localization and Reelin signal transduction. J Biol Chem 280:9671–9677.

Trommsdorff M, Gotthardt M, Hiesberger T, Shelton J, Stockinger W, Nimpf J, Hammer RE, Richardson JA, Herz J (1999) Reeler/Disabled-like disruption of neuronal migration in knockout mice lacking the VLDL receptor and ApoE receptor 2. Cell 97:689–701.

Trotter J, Lee GH, Kazdoba TM, Crowell B, Domogauer J, Mahoney HM, Franco SJ, Muller U, Weeber EJ, D’Arcangelo G (2013) Dab1 is required for synaptic plasticity and associative learning. J Neurosci 33:15652–15668.

Ventruti A, Kazdoba TM, Niu S, D’Arcangelo G (2011) Reelin deficiency causes specific defects in the molecular composition of the synapses in the adult brain. Neuroscience 189:32–42.

Visel A, Thaller C, Eichele G (2004) GenePaint.org: an atlas of gene expression patterns in the mouse embryo. Nucleic Acids Res 32:D552–556.

Wyeth MS, Pelkey KA, Yuan X, Vargish G, Johnston AD, Hunt S, Fang C, Abebe D, Mahadevan V, Fisahn A, Salter MW, McInnes RR, Chittajallu R, McBain CJ (2017) Neto Auxiliary Subunits Regulate Interneuron Somatodendritic and Presynaptic Kainate Receptors to Control Network Inhibition. Cell Rep 20:2156–2168.

